# Group-Specific Carbon Fixation Activity Measurements Reveal Distinct Responses to Oxygen Among Hydrothermal vent *Campylobacteria*

**DOI:** 10.1101/2020.11.29.402834

**Authors:** Jesse McNichol, Stefan Dyksma, Marc Mußmann, Jeffrey S. Seewald, Sean P. Sylva, Stefan M. Sievert

**Affiliations:** Department of Biological Sciences, University of Southern California; Faculty of Technology, Microbiology - Biotechnology, University of Applied Sciences Emden/Leer; Division of Microbial Ecology, University of Vienna; Marine Chemistry & Geochemistry, Woods Hole Oceanographic Institution; Biology Department, Woods Hole Oceanographic Institution

## Abstract

Molecular surveys of low temperature deep-sea hydrothermal vent fluids have shown that *Campylobacteria* (prev. *Epsilonproteobacteria*) often dominate the microbial community and that three subgroups - *Arcobacter*, *Sulfurimonas* and *Sulfurovum -* frequently coexist. In this study, we used replicated radiocarbon incubations of deep-sea hydrothermal fluids to investigate the activities of each group under three distinct incubation conditions. In order to quantify group-specific radiocarbon incorporation, we used newly designed oligonucleotide probes for *Arcobacter, Sulfurimonas,* and *Sulfurovum* to quantify their activity using catalyzed-reporter deposition fluorescence in-situ hybridization (CARD-FISH) combined with fluorescence-activated cell sorting. All three groups actively fixed CO_2_ in short-term (~ 20 h) incubations with either nitrate, oxygen, or no additions (control) at similar per-cell carbon fixation rates. Oxygen additions had the largest effect on community composition and overall cell numbers, and caused a pronounced shift in community composition at the amplicon sequence variant (ASV) level after only 20 h of incubation for all three groups. Interestingly, the effect of oxygen on carbon fixation rates appeared to depend on the initial starting community. Higher carbon fixation rates in oxygen-amended treatments were noted for all three taxa after an unintended disturbance to the sample site that may have selected for more oxygen-tolerant phylotypes. When viewed from a coarse taxonomic level, our data support assertions that these chemoautotrophic groups are functionally redundant in terms of their core metabolic capabilities since they were simultaneously active under all incubation conditions. In contrast, the higher resolution of amplicon sequencing allowed us to reveal finer-scale differences in growth that likely reflect adaptation of physiologically-distinct subtypes to varying oxygen concentrations *in situ*. Despite this progress, we still know remarkably little about the factors that maintain genomic diversity and allow for stable co-existence among these three campylobacterial groups. Moving forward, we suggest that more subtle biological factors such as enzyme substrate specificity, motility, cell morphology, and tolerance to environmental stress should be more thoroughly investigated to better understand ecological niche differentiation at deep-sea hydrothermal vents.

## Introduction

In deep-sea hydrothermal vent ecosystems and other sulfidic, low-oxygen environments, one of the most abundant and biogeochemically-important groups is the *Campylobacteria* (previously known as *Epsilonproteobacteria*) (Waite *et al.*, 2017), whose dominance at vents was first reported 25 years ago (Haddad *et al.*, 1995; Moyer *et al.*, 1995; Polz and Cavanaugh, 1995). Environmental surveys in sulfidic cave systems and at hydrothermal vents have consistently shown that *Campylobacteria* occupy a high-sulfide, low-oxygen niche compared to chemoautotrophic *Gammaproteobacteria* (Macalady et al., 2008; Meier et al., 2017), but less is known about factors driving niche differentiation within the *Campylobacteria*. At a broad level, genomic analyses have revealed that the *Campylobacteria* can be divided into two major orders that have clear differences in their metabolic potential and core bioenergetics. The first order - the *Nautiliales* - was the first to be isolated in pure culture from hydrothermal vents and was originally described as moderately thermophilic hydrogen-dependent obligate anaerobes or microaerobes that tolerate or use a small amount of oxygen (Miroshnichenko *et al.*, 2002, 2004), though aerobic representatives have been more recently identified (Winkel *et al.*, 2014). The second order - the *Campylobacterales* - were isolated at a later time from vent systems (Campbell *et al.*, 2001; Takai *et al.*, 2003), though a close relative (*Sulfurimonas denitrificans*) had been unknowingly cultivated from coastal sediments decades prior (Timmer-Ten Hoor, 1975; Muyzer *et al.*, 1995; Takai et al., 2006). In contrast to the *Nautiliales,* most cultured representatives of the *Campylobacterales* thrive at lower temperatures and can typically tolerate a much greater range of oxygen concentrations (Sievert and Vetriani, 2012, and references therein).

In contrast to these broad, inter-order or inter-class differences in physiology, much less is known about differences in metabolism between members of the *Campylobacterales* and factors governing their co-existence in the natural environment. At our study site (*Crab Spa*), both the chemical composition of hydrothermal fluids (Reeves *et al.*, 2014; McNichol *et al.*, 2016) and the microbial population structure has remained relatively stable since observations began in 2007 (Sievert and Gulmann, unpublished). Similar to other diffuse-flow hydrothermal vents, the microbial community of the subseafloor-derived fluid of Crab Spa is primarily dominated by the three campybacterial groups^1^ of *Campylobacterales*: *Arcobacter, Sulfurimonas,* and *Sulfurovum* (Huber et al., 2007; Meyer et al., 2013; Akerman *et al.*, 2013; Gulmann *et al.*, 2015; Han and Perner, 2015; Stokke *et al.*, 2015; Anderson *et al.*, 2017; Meier *et al.*, 2017; McNichol *et al.*, 2018). Their co-existence over nearly a decade in a chemically-stable environment is indirect evidence they are simultaneously active under natural conditions.

Information on the physiology of these organisms primarily comes from pure culture work which has shown that isolates of *Sulfurimonas* and *Sulfurovum* typically use H_2_ and reduced sulfur compounds as electron donors and nitrate, sulfur, and oxygen as electron acceptors (Sievert and Vetriani, 2012 and references therein). Despite this relatively limited repertoire of substrates, isolates differ with respect to the presence/absence of pathways. For example, some isolates are strict microaerobes, whereas others can use nitrate as an alternative electron acceptor. In addition, oxygen tolerance varies markedly (even within the same group), with cultured representatives spanning a spectrum from obligate anaerobe (Mino et al., 2014) to obligate aerobes that can tolerate high levels of oxygen in the presence of high exogenous CO_2_ concentrations (Mori *et al.*, 2018). Environmental surveys and short-term incubations have also underscored the importance of oxygen in structuring natural communities and affecting carbon fixation efficiency (Meier et al., 2017, McNichol et al 2018).

Much less is known about *Arcobacter*, since no chemoautotrophic pure culture isolate exists. Based on enrichment cultures, it has been inferred to be a chemoautotroph possibly thriving under highly turbulent and sulfidic conditions (Sievert et al., 2007; 2008). Modern ‘omics surveys and the genome reconstructions have similarly shown that the same general pathways are present for diverse *Campylobacteria* (Meier et al., 2017; Tully et al., 2018; Galambos *et al.*, 2019) and that they are transcriptionally active (Fortunato and Huber, 2016; Galambos et al., 2019; Fortunato and Huber, 2016). However, few studies have directly measured primary production under conditions reflective of natural conditions (McNichol et al., 2018), much less at a resolution possible to distinguish differences between groups of *Campylobacteria*. Thus, the factors driving the activity and chemoautotrophic carbon fixation in the natural environment remain largely unknown.

To better quantify the activity of these three groups in diffuse-flow hydrothermal fluids under varying conditions, we applied a recently-developed method for the quantification of group-specific radiocarbon incorporation (Dyksma *et al.*, 2016) in combination with newly designed oligonucleotide probes to quantify group-specific carbon fixation activity under three distinct and replicated experimental conditions. Combined with concurrently obtained chemical measurements and molecular data, this work provides new insights into the activity and ecophysiology of *Campylobacterales* at deep-sea hydrothermal vents.

## Materials and methods

### Hydrothermal fluid sampling and paired/radiocarbon incubations

Samples were obtained from the low-temperature hydrothermal vent *Crab Spa,* which has been described in detail elsewhere (Reeves *et al.*, 2014; McNichol *et al.*, 2016, 2018). Fluid samples were obtained from *Crab Spa* using Isobaric Gas-Tight Samplers [IGTs; (Seewald *et al.*, 2002)]. The use of radiocarbon for the incubations to measure carbon fixation made it logistically impossible to conduct them under *in situ* pressure in IGTs due to an unacceptable risk of ^14^C contamination outside of the ship’s radioisotope van. These experiments are referred to as ‘radiocarbon incubations’ in the subsequent text. Synchronized non-radioactive incubations (with the same source fluids and headspace:liquid ratio) were also carried out to measure additional chemical and biological parameters (cell counts, CARD-FISH, 16S rRNA gene amplicon sequencing) and the concentration of potential electron donors / acceptors and metabolic by-products over the incubation period. These are referred to as ‘paired incubations’ below.

For radiocarbon incubations, 8 mL of hydrothermal fluid was transferred from the IGTs into 12 mL N_2_-flushed exetainer screw cap vials (Labco Limited) and spiked with ^14^C sodium bicarbonate after transport to the radioisotope van (specific activity 56 mCi mmol^−1^, Hartmann Analytic, Germany) resulting in a final radioactive DIC fraction of 0.17%. All vials were incubated for 16 h at *in situ* temperature (24 °C) in the dark after which cells were fixed with formaldehyde (2%, final concentration) for 1 h at room temperature. For paired incubations, approximately 80 mL of hydrothermal fluid was transferred with a sterile needle directly from the IGT into a N_2_-purged 123 mL GL45 Pyrex flask sealed with a ~2cm thick butyl stopper (Part #444704, Glasgerätebau Ochs, Germany), and excess N_2_ overpressure released with another needle at the end of filling. Flasks were rinsed with MilliQ water between incubations, and sterilized by contact with 70% ethanol and subsequent drying in a 50 °C oven. In both radiocarbon and paired incubations, nitrate was added as a small aliquot of a N_2_-purged 0.175 M NaNO_3_ solution, and O_2_ was added as gas directly to the headspace to reach an initial concentration of dissolved O_2_ of approximately 100 μM. Concentrations of dissolved O_2_ were calculated as described previously (Weiss, 1970).

Cell counts and chemical measurements (only for paired incubations) were conducted as previously described (McNichol *et al.*, 2016) except oxygen was measured with the Fibox 3 instrument using an optode spot affixed to the inside of the bottle using silicone (Pts3, PreSens, Regensburg, Germany). Subsamples were taken with sterile needles/syringes that were purged with 0.2 μm-filtered N_2_ gas three times prior to sampling. The same volume of sterile N_2_ gas as the fluid sample was injected to prevent underpressure forming in the incubation vessels.

### CARD-FISH and sample preparation for flow cytometry

Samples from radiocarbon incubations were filtered onto 25 mm polycarbonate membrane filters with a 0.2 μm pore size (GTTP, Millipore, Eschborn, Germany). Permeabilization and CARD-FISH was performed as described previously (Pernthaler *et al.*, 2002) with the following modifications. Samples for flow cytometry were not embedded in agarose. Endogenous peroxidases were inactivated in 3% H_2_O_2_ for 10 min at room temperature. The hybridization temperature was 46°C and washing was performed at 48°C according to the protocol of (Ishii *et al.*, 2004). An overview of oligonucleotide probes used in this study is shown in Supplementary Table S1. Tyramides labelled with Alexa Fluor 488 fluorescent dye (Molecular Probes, USA) were used for CARD signal amplification. Cells were removed from filter membranes using a cell scraper or were vortexed in 5 ml of 150 mM NaCl containing 0.05% Tween 80 according to (Sekar *et al.*, 2004). Prior to flow cytometry, large suspended particles were removed by filtration through 8 μm pore-size filter (Sartorius, Göttingen, Germany) to avoid clogging of the flow cytometer.

### Fluorescence activated cell sorting (FACS) and scintillation counting

Flow sorting of CARD-FISH-identified cells and scintillation counting of sorted cell fractions were performed as described previously (Dyksma *et al.*, 2016). In brief, cells were sorted using a FACSCalibur flow cytometer (Becton Dickinson, Oxford, UK) in batches of at least 10,000 cells. Hybridized cells were identified on scatter dot plots of green fluorescence versus 90° light scatter and background was determined by flow cytometric analysis of hybridizations with a nonsense probe (NON338) prior to flow sorting. Sorted cell batches that were collected on polycarbonate filters were transferred into 5 ml scintillation vials and mixed with 5 ml UltimaGold XR (Perkin Elmer, Boston, USA) scintillation cocktail. Radioactivity of sorted cell batches was measured in a liquid scintillation counter (Tri-Carb 2900, Perkin Elmer, USA). Average cell-specific carbon fixation rates were calculated according to the equation *R* = (*A*\**M*)/(*a*\**n*\**t*\**L*). *A* represents the radioactivity of the sorted cell batch in Becquerel (Bq), *M* represents the molar mass of carbon (g/mol), *a* represents the specific activity of the tracer (Bq/mol), *n* represents the number of sorted cells, *t* represents the incubation time and *L* equals to the ratio of total DIC to ^14^C-DIC. Readers should note that this population cell-specific average represents a slightly different calculation compared to studies that measure single-cell incorporation and average across all individually-measured cell uptake rates. Another key difference of this technique versus single-cell assays is that tens of thousands of cells can be sorted and quantified precisely versus dozens to hundreds for techniques such as NanoSIMS.

### CARD-FISH probe design and testing

To demarcate the regions of the SILVA reference tree (version: “SILVA_119_SSURef_Nr99”) to target for probe design, steps from the ARB tutorial by Amy Apprill (http://www.arb-home.de/documentation.html) were used as a template. Briefly, partial 16S genes derived from PCR amplicons sequenced with 454 technology [from the Atlantis cruise AT26-10 (McNichol *et al.*, 2018)] were first aligned with the SINA web aligner (Pruesse *et al.*, 2012). Clusters without partial 16S sequences derived from AT26-10 were removed (e.g., those containing *Sulfurimonas denitrificans* and *Sulfurimonas gotlandica* that occur in non-vent environments) and the remaining sequences were used for probe design.

Candidate probe sequences were selected based on coverage of the relevant groups (*Arcobacter*, *Sulfurimonas,* and *Sulfurovum*), and helper/competitor probes were also designed to assist with specific binding (Supplementary Table S1). Probes were tested *in silico* against full-length clone sequences derived from the same vent field (Gulmann *et al.*, 2015) and also (where applicable) partial 16S SSU rRNA amplicon sequences derived from single-amplified genomes sorted by the Bigelow Single-Cell Genomics Center (unpublished data). We note that although probes already existed for *Arcobacter* (ARC94 & ARC1430: Snaidr et al., 1997; Wirsen et al., 2002), they only cover about 80% of *Arcobacter* sequences found in SILVA SSU database r138 (Quast *et al.*, 2013). In addition, ARC94 has unacceptable outgroup matches to *Nitratiruptoraceae* (67% coverage) and *Thioreductoraceae* (100% coverage), both members of the *Nautiliales* which have markedly different physiologies compared to *Arcobacter* and have been shown to be present alongside *Sulfurimonas* at least under some conditions (McNichol *et al.*, 2018). In contrast, our newly designed *Arcobac*ter probe (ARC673) increases group coverage to ~ 94% and has no significant outgroup matches.

To determine the optimum formamide concentration for hybridization, probes (with helpers and competitor oligonucleotides) were tested across a formamide melting curve with positive controls and 1-mismatches as described in Supplementary Table S1. To ensure that probes only hybridized with *Campylobacteria* and did not cross-hybridize between groups or to non-campylobacterial cells, a series of double-hybridizations were carried out with each of the new probes in combination with either EPSI549/914 or each other. These were carried out on background (unincubated) samples from *Crab Spa*. Images of these double-hybridizations and CARD-FISH counts from experiments are available online at https://osf.io/yts4p/.

### 16S rRNA sequence analysis

DNA extraction and sequencing was as described in McNichol et al (2018) for low-volume extractions of biomass captured on a 0.2 μm Sterivex filter and subsequent 454 amplicon sequencing with the 27Fmod/519Rmodbio primers. Raw fastq files were manually de-multiplexed from SFFs file using qiime1 (Caporaso *et al.*, 2010), and imported into qiime2 (2018.8 build; Bolyen *et al.*, 2018). Sequences were denoised using the q2-DADA2 plugin (Callahan *et al.*, 2016), and classified using the naive Bayesian classifier (Bokulich *et al.*, 2018) in qiime2 using SILVA 132 as a reference database (Quast *et al.*, 2013) subsetted to the primer region. All scripts and accessory files needed to reproduce this analysis are available at: https://osf.io/9e6ra/. NMDS plots were made with PAST (Hammer *et al.*, 2001) using an ASV table subsetted to include only *Campylobacteria*. Raw fastq files split by sample were deposited at the NCBI SRA under Bioproject number PRJNA505918.

## Results and Discussion

Compared to previously reported incubations of hydrothermal vent fluids that mainly focused on biogeochemical rate measurements (Tuttle *et al.*, 1983; Wirsen *et al.*, 1986, 1993; Perner *et al.*, 2010, 2011, 2013; Bourbonnais *et al.*, 2012; McNichol et al., 2018; Böhnke *et al.*, 2019; Trembath-Reichert *et al.*, 2019), our experiments provide finer-scale assessment of microbial community activity and document changes in taxonomic composition during incubations in greater detail. This was possible because we simultaneously measured changes in fluid chemistry, microbial carbon fixation, overall cell growth, and changes in microbial community composition. Of particular note is the design and application of novel olignucleotide probes for *Sulfurimonas*, *Sulfurovum* and *Arcobacter* which increased probe coverage and specificity versus existing probes (Methods; Supplementary Table S1). When combined with a novel technique that uses ^14^C-DIC tracer incubations and flow cytometric sorting (Dyksma *et al.*, 2016), this allowed us to specifically quantify carbon fixation among distinct campylobacterial populations.

We also took care to minimize sampling artifacts by beginning incubations as quickly as possible after retrieval from the submersible (on average ~7 hours after sampling at the seafloor; Table 1). To simulate three different redox conditions, we either made no additions or provided nitrate / oxygen as amendments. We chose to add electron acceptors rather than donors because the fluid chemistry of *Crab Spa* is characterized by high sulfide concentrations (~180μM ΣH_2_S; Reeves *et al.*, 2014) relative to oxygen and nitrate (O_2_ ~3 μM; NO_3_^−^ ~6 μM), leading to electron acceptor limitation of microbial growth (McNichol *et al.*, 2016).

**Table 1.** Summary of conditions during atmospheric pressure incubations using IGT fluid samplers

| Amendment* | Concentration<br>(μM) | Temperature<br>($^{\circ}$C) | # Replicates | Time from seafloor*<br>(h) | Experimental IDs<br>(Alvin dive #s, IGT ID) |
| --- | --- | --- | --- | --- | --- |
| Control | NA | 24 | 2 | 7.3, 5.1 | 4772-8, 4773-7 |
| Nitrate | 100-200 | 24 | 3 | 9.3, 6.2, 8.4 | 4769-4, 4770-8, 4773-5 |
| Oxygen | 110 | 24 | 3 | 4.6, 9.3, 7.0 | 4770-5, 4772-4, 4773-8 |

\*Time from sampling at seafloor to beginning of incubation

### General observations

Based on all sorted *Campylobacteria* (EPSI 549/914-hybridized), the average cell-specific carbon fixation rates ranged from 13 to 36.3 fg C cell^−1^ day^−1^ (average = 23.6 ± 8.5; n=8; Fig 1), which were approximately one third of that previously measured by NanoSIMS in incubations conducted at *in situ* pressure with isobaric gas-tight samplers which underwent a similar CARD-FISH hybridization procedure (25.4 - 134.2, average = 78.4 ± 25.3 fg C cell^−1^ day^−1^ (n=18); McNichol *et al.*, 2018), possibly indicating a pressure effect. All three targeted groups demonstrated radiocarbon incorporation at comparable rates, however, their activity was highly variable between the replicates and amendments (*Arcobacter* 36.8 ± 26.9; *Sulfurimonas* 37.5 ± 24.7; *Sulfurovum* 38.8 ± 29.0 fg C cell^−1^ day^−1^).

**Figure 1:**
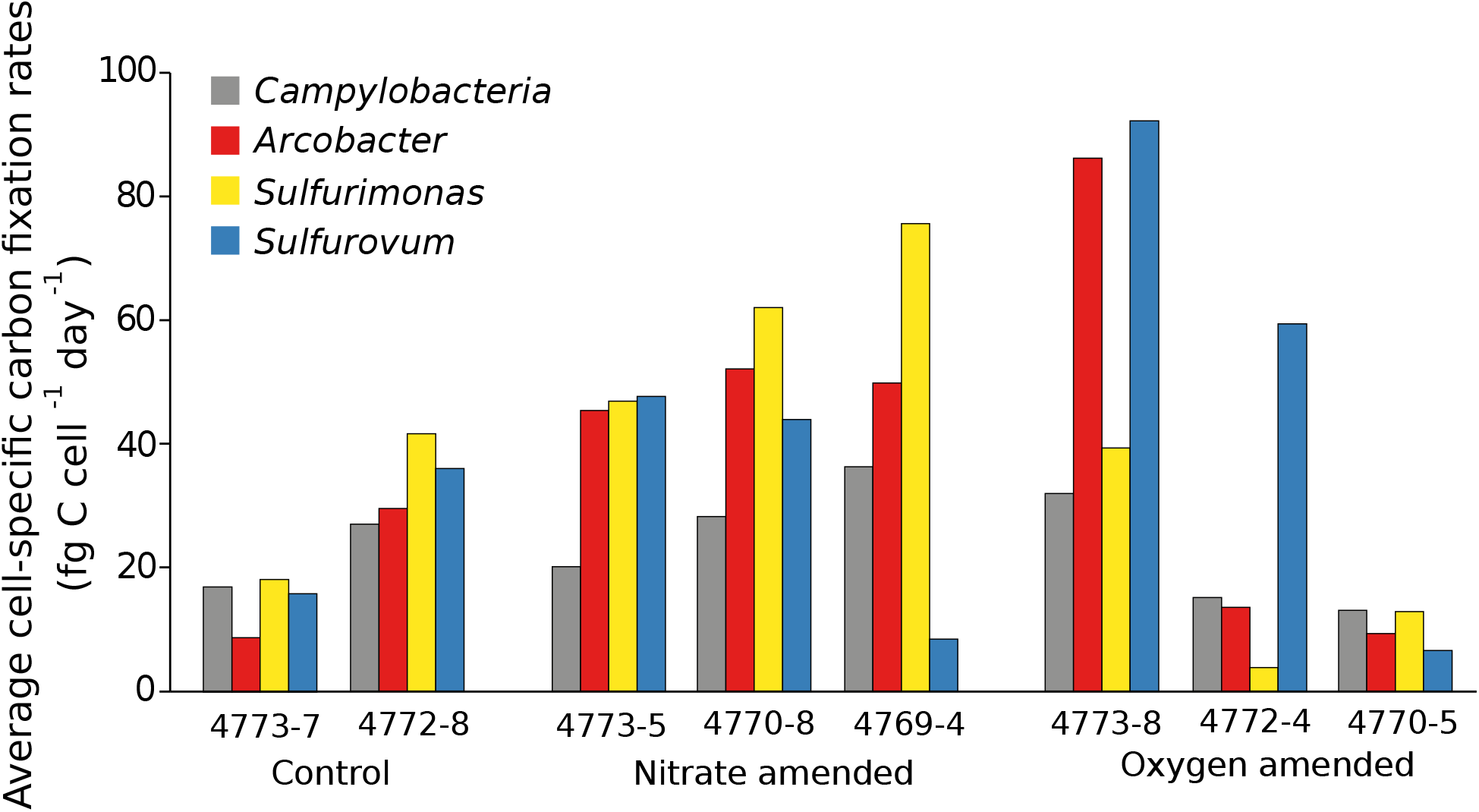
Results of group-specific radiocarbon incorporation during short-term microbial incubations at atmospheric pressure.

Similar to previously published data (McNichol *et al.*, 2016), active microbial metabolism in the paired incubations was apparent based on the rapid consumption of sulfide, nitrate and oxygen at similar bulk rates. Control incubations (no additions) were limited by electron acceptors as demonstrated by the cessation of sulfide consumption after oxygen and nitrate (in that order) were successively drawn down (Fig 2a, left panel). In contrast, in oxygen/nitrate addition experiments where electron acceptors were not limiting, sulfide was continuously consumed (Fig 2a, middle and right panel). Determining reaction stoichiometry by making comparisons of nitrate/oxygen consumption as described in McNichol *et al.* (2016) is not straightforward for these incubations because oxygen was supplied from the headspace and measured in the fluids. Thus, oxygen drawdown as shown in Fig 2a likely underestimates the total amount of oxygen consumed. The fraction of nitrate likely processed through the dissimilatory nitrate reduction to ammonium (DNRA) pathway varied considerably between the three replicates to which nitrate was added (4769-4 = 48%, 4770-8 = 12%, 4773-5 = 31%; values are corrected to account for ~15 fg ammonium uptake per new cell), with the remainder presumably being denitrified to N_2_.

**Figure 2:**
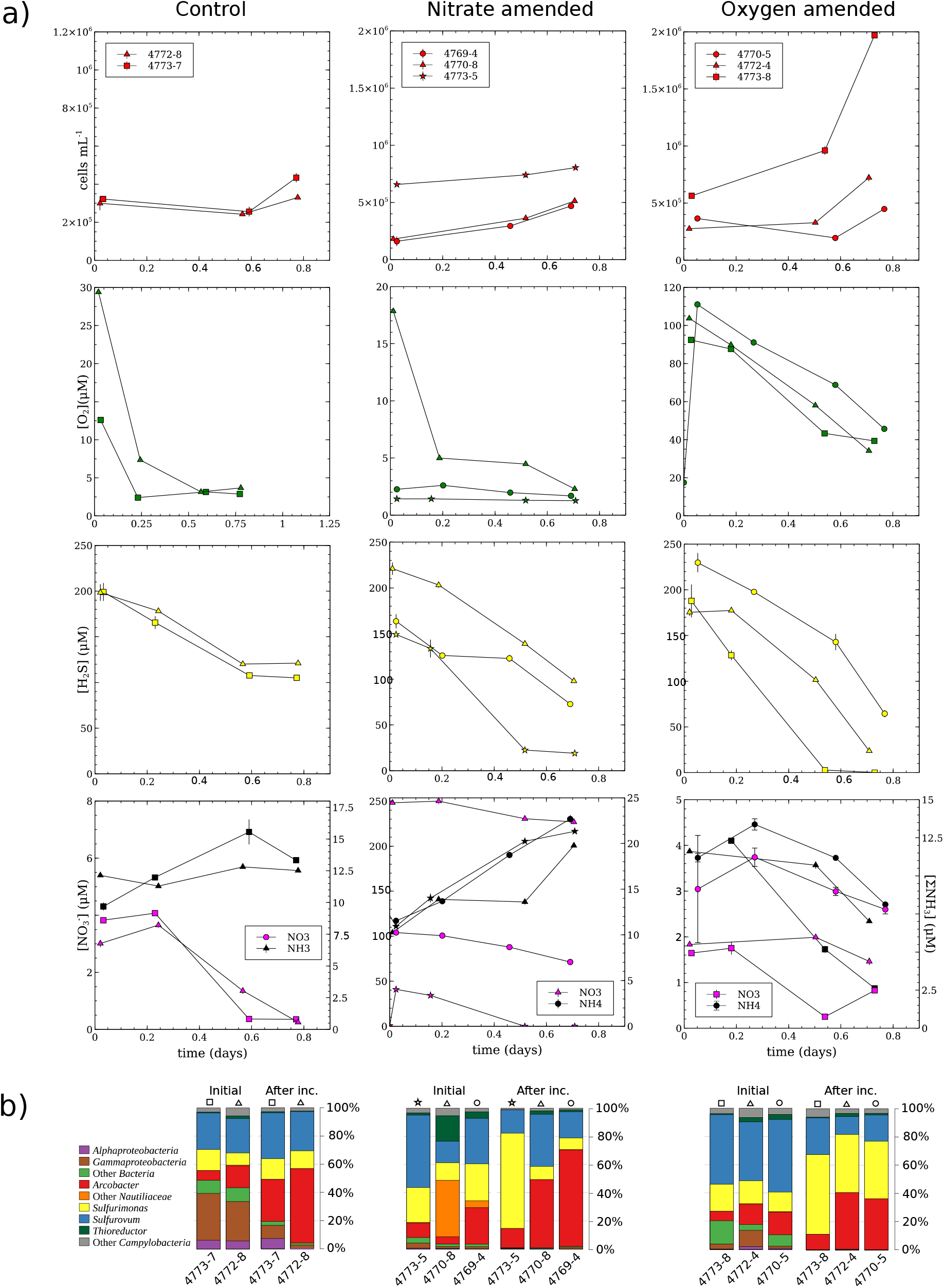
Paired incubation growth dynamics and community composition changes.

### General community patterns during incubations as inferred by 16S amplicon analysis

In addition to large changes in concentrations of electron donors/acceptors and increases in cell counts, 16S rRNA gene amplicon sequence analysis from the paired incubations showed clear evidence of growth during the incubation period. The initial community was dominated by *Campylobacterales*, though some samples contained a significant fraction of likely heterotrophic *Proteobacteria* (up to ~ 40 %; Fig 2b) and one sample (4770-8) had a single ASV from the *Nautiliales* that accounted for 27% of the total relative abundance at the beginning but was absent at the end of incubations. Despite providing three different conditions including one which was consistently oxic, amplicons at the end of incubations were all dominated by *Campylobacteria.* The gammaproteobacterial SUP05 clade which thrives in hydrothermal fluids with a higher proportion of oxic seawater (Meier *et al.*, 2017) was a minor component of overall sequence counts at the beginning and end of incubations (≤1%), suggesting they were not important chemoautotrophs *in situ* at *Crab Spa* or during the incubations. In addition, the fraction of putatively heterotrophic bacteria had decreased to less than 5 % by the end of incubations in all but one case, suggesting these organisms also did not grow during incubations (Fig 2b).

At the end of the incubations, CARD-FISH and amplicon sequences (ASVs summed within each group) showed a relatively even abundance of the three target groups *Sulfurimonas, Sulfurovum* and *Arcobacter* (Fig 2b, 3, S1). Interestingly, this was not the case in high pressure incubations conducted earlier in the same year at the same study site. In those incubations, amplicon sequences and CARD-FISH cell counts showed that *Sulfurimonas* (and to a lesser extent *Thioreductor,* a member of the *Nautiliales*) dominated activity and biomass, whereas *Arcobacter* and *Sulfurovum* were present at low abundance at the end of incubations (McNichol *et al.*, 2018). These *in situ* pressure incubations did not measure the starting community composition as we reported here, but given that *Arcobacter* and *Sulfurovum* are always present in initial samples (Fig 2b) it appears likely that the robust growth of these taxa in our incubations at atmospheric pressure is an effect specific to our experimental setup (e.g. perhaps due to the presence of a headspace gas phase as discussed further below). These differing incubation conditions might also explain the lack of growth of *Thioreductor* in this study, but we note that the abundance of related organisms is more variable (see for e.g. *Nautiliales* in Fig 2b 4770-8, initial time point), meaning that the effect of initial community composition might be more important for growth of these putatively thermophilic taxa in incubations. In this regard, it is worth noting that *Sulfurimonas* accounted for significantly higher proportion of the natural community in January 2014 compared to November 2014 (Sievert and Gulmann, unpublished data), possibly also contributing to the dominance of *Sulfurimonas* in the IGT incubations reported by McNichol et al. (2018).

**Figure 3:**
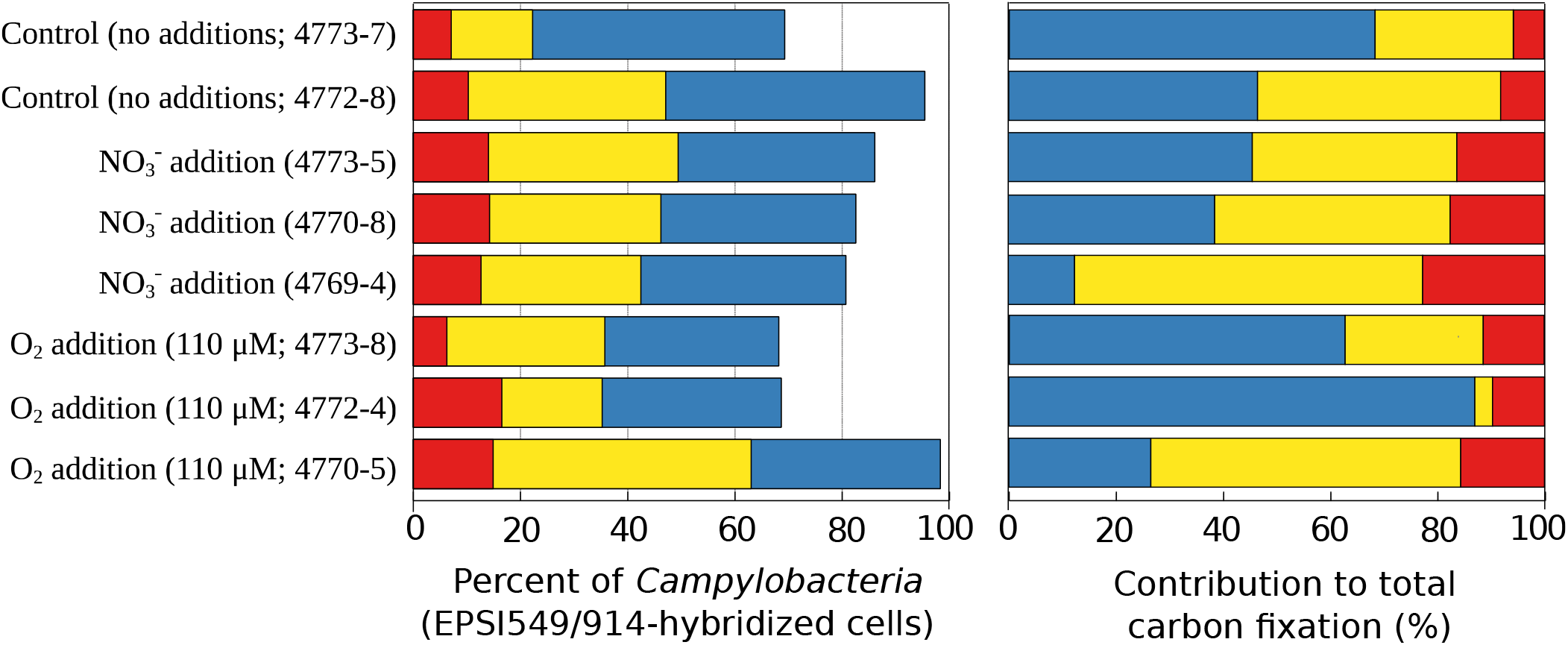
Percentages of total cells identified in *radiocarbon incubations* using CARD-FISH and their predicted relative contributions to carbon fixation within *Campylobacteria* based on cell-specific rate measurements.

### Chemoautotrophic rates and community composition were most influenced by oxygen

Despite similar activities at the group level, ASV analysis revealed that the final community compositions in oxygen amendments diverged markedly from control and nitrate additions in a consistent manner in samples taken across all dives (Fig 4 and Supplementary Fig S2). This supports the notion that oxygen concentration is a major physicochemical factor determining community structure *in situ,* in line with observations from environmental surveys and incubation studies (Meier *et al.*, 2017; McNichol *et al.*, 2018). A clear “bottle effect” could also be observed, with the composition diverging markedly from the initial state even for no-amendment controls (Fig 4).

**Figure 4:**
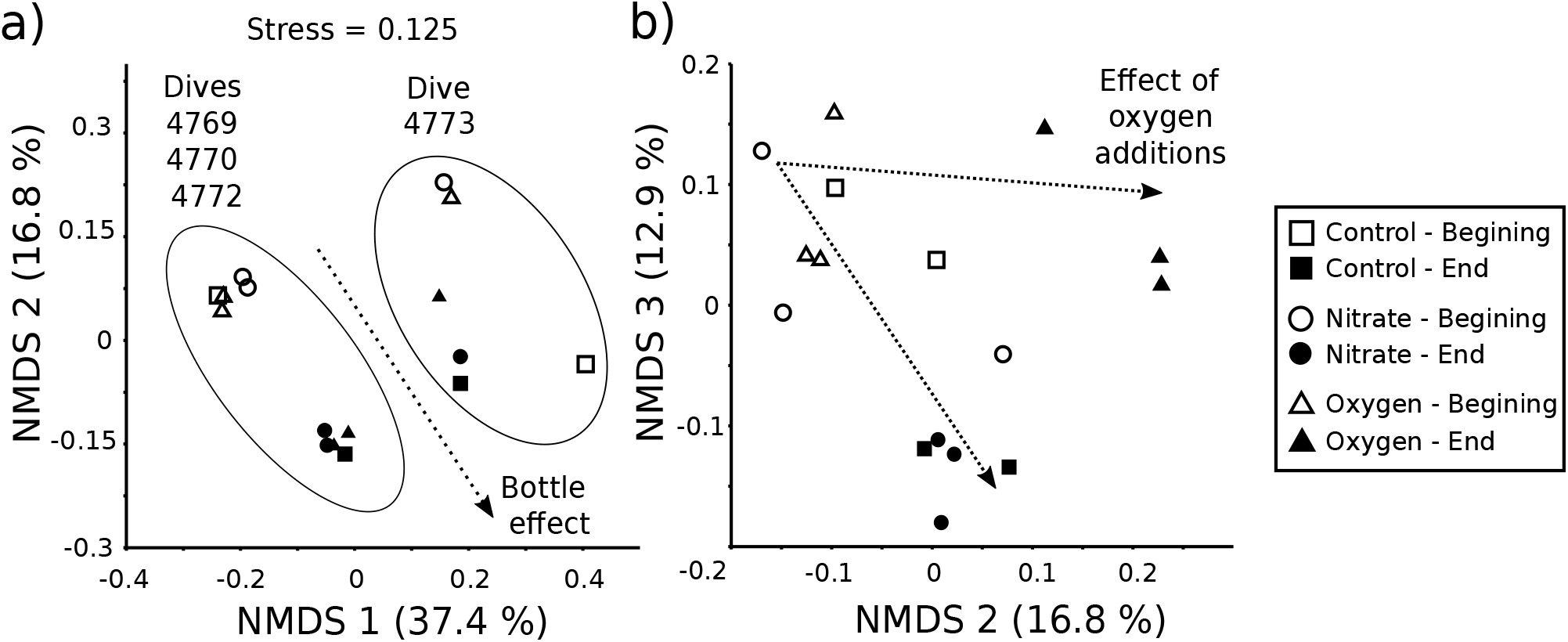
Non-metric multidimensional scaling (NMDS) plots showing changes in campylobacterial community composition across incubations and sampling time. Panel a) shows the major community difference between *Alvin* dive 4773 and other dives which is likely attributed to the deployment of an *in-situ* sampling device (discussed in text), as well as a clear “bottle effect” for all samples. Panel b) shows a reproducible effect of oxygen on community composition regardless of dive.

Interestingly, the effect of oxygen on community carbon fixation appeared to be strongly linked to the initial community composition in the sampled fluids. Although the microbial community sampled at *Crab Spa* was fairly consistent across the first three Alvin dives, we observed a large difference in campylobacterial community composition after the deployment of an instrument at *Crab Spa* at the end of dive 4772 (after samples for these incubations were taken; Fig 4). Strikingly, there was essentially no overlap in ASVs among the initial communities before and after instrument deployment, suggesting that a large and rapid (~24 h) community shift occurred at *Crab Spa* at a fine-scale taxonomic level. Cell counts from incubations 4773-5 and 4773-8 also had considerably higher starting cell densities, and rapid cell growth and drawdown of oxygen was observed for both (Fig 2a, right panel). In addition, we observed much higher cell-specific carbon fixation rates for all three campylobacterial groups in the oxic incubation (4773-8), with the difference being most notable for

*Arcobacter* and *Sulfurimonas*. In contrast, cell-specific carbon fixation rates for nitrate/control incubations were much more similar to incubations with fluids obtained in the previous dives. Based on this, we hypothesize that taxa present in fluids obtained on dive 4773 may have been more tolerant to oxygen due to the unintended disturbance to the natural system described above. This hypothesis is in line with knowledge from culture work that has shown that optimum oxygen concentrations for *Campylobacteria* can be as low as 1-10 μM (Sievert *et al.*, 2007) or approach nearly full oxygen saturation if carried out in sealed vessels with exogenous CO_2_ (Inagaki et al., 2004; Takai et al., 2006; Mori et al., 2018). Overall, this indicates that previous observations of lower growth efficiency for *Campylobacteria* under oxic conditions (McNichol et al., 2018) may not be easily generalized since they would depend on the community composition of the *in situ* microbial community.

### Evidence of high chemoautotrophic activity and the ability to respond rapidly to changing conditions for *Arcobacter*

Chemoautotrophic *Arcobacter* have been characterized as microaerophilic sulfur-oxidizing bacteria that can tolerate high sulfide and produce copious filamentous sulfur (Taylor and Wirsen, 1997; Wirsen *et al.*, 2002). The one characterized chemoautotrophic enrichment culture (*Candidatus* Arcobacter sulfidicus) is highly motile, and forms aggregates at the oxic-anoxic interface (Sievert *et al.*, 2007). Similar organisms have been suggested to be important players in highly turbulent environments with high sulfide concentrations, such as those found after a volcanic eruption (Sievert *et al.*, 2008; Meyer et al., 2013). In our incubations, we observed evidence of growth and carbon fixation for *Arcobacter*-affiliated organisms. The relative abundance of 16S rRNA gene amplicons affiliating with *Arcobacter* increased in most incubations independent of the treatment (Fig 2b), indicating cell growth. Supporting this, only a few *Arcobacter* ASVs were dominant at the end of incubations compared to the beginning and were completely distinct between oxygen-amended and the other incubations (Supplementary Fig S2a). This suggests quick growth of a relatively small number of phylotypes within this group that are adapted to different oxygen concentrations *in situ*.

Strikingly, one of the highest cell-specific carbon fixation rate was determined for *Arcobacter* in the incubation 4773-8 (86.2 fg C cell^−1^ d^−1^; Fig 1). In this incubation, one particular ASV became abundant that was not observed in other incubations (Supplementary Fig S2a). The *Arcobacter* average community cell-specific rate is considerably higher than the value of 14.4 fg C cell^−1^ d^−1^ previously reported for *Candidatus* Arcobacter sulfidicus (Wirsen et al., 2002), which indicates these organisms were growing well under our incubation conditions. Although the average cell-specific carbon fixation rates for *Arcobacter* were comparable to those of *Sulfurimonas* and *Sulfurovum* (*Arcobacter*, 36.8 fg C cell^−1^ d^−1^; *Sulfurimonas*, 37.5 fg C cell^−1^ d^−1^; *Sulfurovum*, 38.8 fg C cell^−1^ d^−1^), they only contributed 5.7-23% to total carbon fixed due to their lower overall cell counts (Fig 3).

Despite lower cell abundances, *Arcobacter* represented a larger fraction of total amplicon sequences and were more likely to be composed of only one or a few ASVs compared to *Sulfurimonas* and *Sulfurovum* (Supplementary Fig S2a-c). Considering the ratio of ASV abundance to CARD-FISH cell counts, we found that the ratio for *Arcobacter* was the highest and most variable (2.1 ± 1.3), whereas the ratio for both *Sulfurimonas* and *Sulfurovum* was close to 1:1 on average (1.2 ± 0.5 and 1.0 ± 0.1, respectively). This observation indicates *Arcobacter* has a higher average 16S rRNA gene copy number per genome compared to *Sulfurimonas* and *Sulfurovum*, which has been shown to be a proxy for bacterial adaption to resource availability (Roller *et al.*, 2016). Together, these data support the previous speculation that *Arcobacter* can rapidly respond to changing conditions (Sievert *et al.*, 2007, 2008), though it seems that at least some members of all three groups have the same potential to respond rapidly (Supplementary Fig S2b-c).

Pure culture work has shown that some *Arcobacter* strains are capable of true aerobic growth (Vandamme *et al.*, 1991), in contrast to other (heterotrophic) members of the *Campylobacteria* which require microaerophilic conditions and/or exogenous CO_2_ to support their growth (Bolton and Coates, 1983; Al-Haideri *et al.*, 2016). This, combined with the fact that (presumably heterotrophic) *Arcobacter* have been recently identified in sinking sediment particles (Boeuf *et al.*, 2019), could indicate a predisposition of this group for oxic environments where fast growth is favoured. In our incubations, we provided a headspace of nitrogen gas which would have reduced overall CO_2_ concentrations due to outgassing which may have provided microenvironments more suitable for *Arcobacter*, potentially explaining their apparent lack of growth in incubations which did not contain a headspace (McNichol *et al.*, 2018).

### *Sulfurimonas* and *Sulfurovum* account for the majority of carbon fixation

*Sulfurimonas* and *Sulfurovum* accounted for the majority of carbon fixation due to their high cell abundances in the incubations (Fig 3), in line with the finding that they are commonly identified as the dominant microorganisms in diffuse-flow hydrothermal fluids (Huber et al., 2007; Meyer et al., 2013; Akerman *et al.*, 2013; Gulmann *et al.*, 2015; Han and Perner, 2015; Stokke *et al.*, 2015; Anderson *et al.*, 2017; Meier *et al.*, 2017; McNichol *et al.*, 2018). Compared to *Arcobacter,* the contribution of *Sulfurimonas and Sulfurovum* to total carbon fixation was higher, but more variable, accounting for 39.5 ± 21.7 % and 47.8 ± 25.1 %, respectively (Fig 3). During the incubations, both *Sulfurimonas* and *Sulfurovum* mostly maintained a relatively high ASV diversity, but an effect of incubation could still be observed with different ASVs coming to dominate at the end of incubations (in particular for incubation 4773-5; Supplementary Fig S2b-c). Average cell-specific carbon fixation rates for *Sulfurimonas* were consistently high with nitrate additions (61.5 ± 14.4 fg C cell^−1^ d^−1^) compared to oxygen additions (18.6 ± 18.5 fg C cell^−1^ d^−1^). We observed the highest cell-specific carbon fixation rates for *Sulfurovum* in oxygen amended incubations (92.2 fg C cell^−1^ d^−1^; Fig 1), and note that these rates were substantially higher compared to *Sulfurimonas*, with the exception of 4770-5 which showed uniformly low rates for all three groups. This presumably aerobic metabolism is consistent with a report showing positive correlations between oxygen and the abundance of *Sulfurovum*-related sequences in natural systems (Meier *et al.*, 2017). As discussed above, differences in cell-specific carbon fixation rates for *Sulfurimonas / Sulfurovum* in 4772-4 / 4770-5 versus 4773-8 may be related to the differing oxygen tolerances of the initial communities.

## Conclusions

In this study, we obtained group-specific chemoautotrophic carbon fixation rates for *Arcobacter*, *Sulfurimonas,* and *Sulfurovum*, the three dominant chemoautotrophic groups commonly observed in diffuse-flow hydrothermal vent fluids and other sulfidic environments. We also observed that all three groups were simultaneously active in our incubations, that oxygen had the largest effect on microbial community composition on short time-scales, and that the effect of oxygen on carbon fixation rates/efficiency is likely dependent on the specific community members present. This demonstrates that despite our study site’s chemical stability over time, it retains community members that can respond rapidly to changing conditions - most likely due to adaptations to tolerate differing *in situ* oxygen levels. This observation is consistent with data showing that *Sulfurovum* and *Sulfurimonas* are much more diverse at a genomic level compared to gammaproteobacterial sulfur oxidizers of the SUP05 clade found in more oxic seawater surrounding hydrothermal vents (Meier *et al.*, 2017). This greater campylobacterial diversity has been suggested to be due the wide range of micro-niches made available during mixing of hydrothermal fluids with seawater (Meier *et al.*, 2017).

In the future, more careful investigation of precise environmental conditions, environmental genomic diversity, and concerted isolation campaigns could help reveal further reasons for the co-existence of these three taxa and maintenance of high microdiversity among chemoautotrophic *Campylobacteria.* It is likely that previously implied differences in enzyme substrate specificity (Meier *et al.*, 2017; Götz *et al.*, 2019) and the ability to move to physiologically optimal microenvironments (Sievert *et al.*, 2007) are extremely important factors. In addition, more subtle and less well-explored features may include size (Stokke *et al.*, 2015) and arrangement of cells (Mino *et al.*, 2014) which would likely have important implications for attachment to solid substrates, the ability to compete for limiting nutrients, or avoiding grazing/viral lysis.

## Acknowledgements

We thank the officers, crew, and pilots of the R/V Atlantis and HOV Alvin for their expert help at sea and their outstanding efforts acquiring the samples for this study. Special thanks are due to Florian Götz for assistance with shipboard incubations and sulfide measurements. We thank Jörg Wulf for synthesizing the tyramides needed for CARD-FISH, and Niculina Musat for advice designing probes and capturing the CARD-FISH images used in this study. Thanks are also due to Ben van Mooy for lending his laboratory’s oxygen optode system as well as to Scott Wankel, Carly Buchwald, and Zoe Sandwith for help quantifying nitrate/nitrite. This research was funded by the US National Science Foundation grants OCE-1131095 (S.M.S) and OCE-1136727 (S.M.S., J.S.S.). Further support was provided by the WHOI Investment in Science Fund (S.M.S.). Funding for J.M. was further provided by doctoral fellowships from the Natural Sciences and Engineering Research Council of Canada (PGSD3-430487-2013, PGSM-405117-2011) and the National Aeronautics and Space Administration Earth Systems Science Fellowship (PLANET14F-0075), an award from the Canadian Meteorological and Oceanographic Society, and the WHOI Academic Programs Office.

---

**Table S2:**
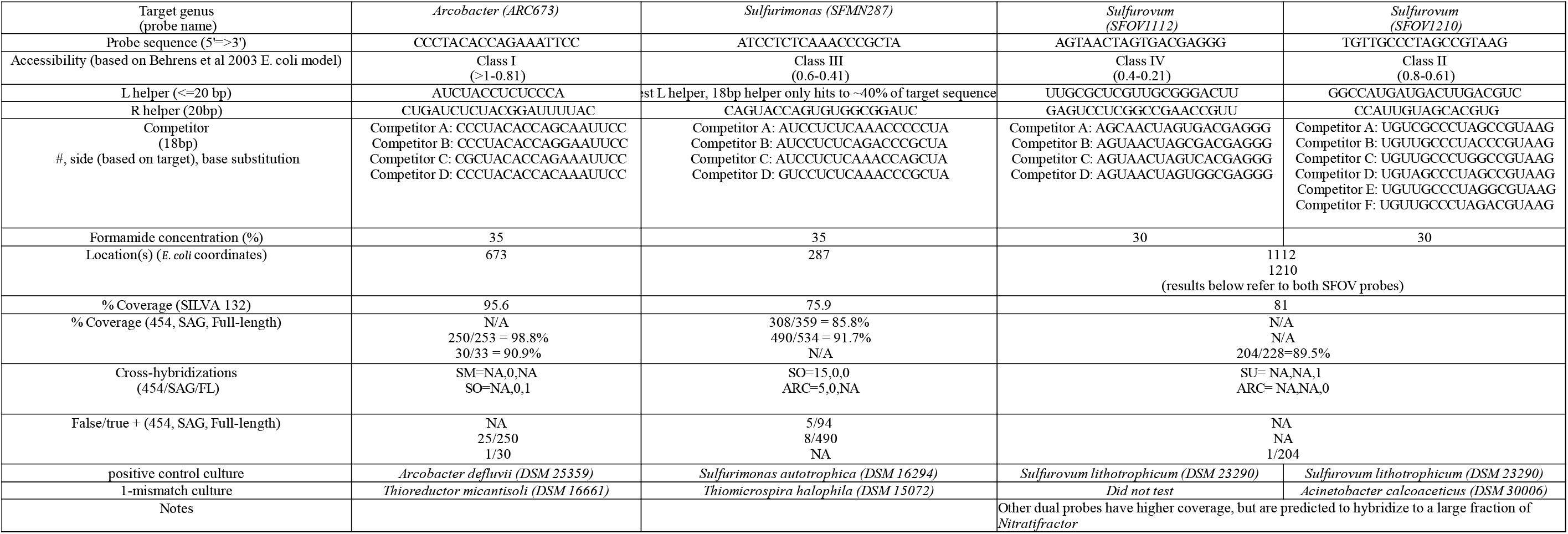
CARD-FISH Probes and Sequences.

**Fig S1:**
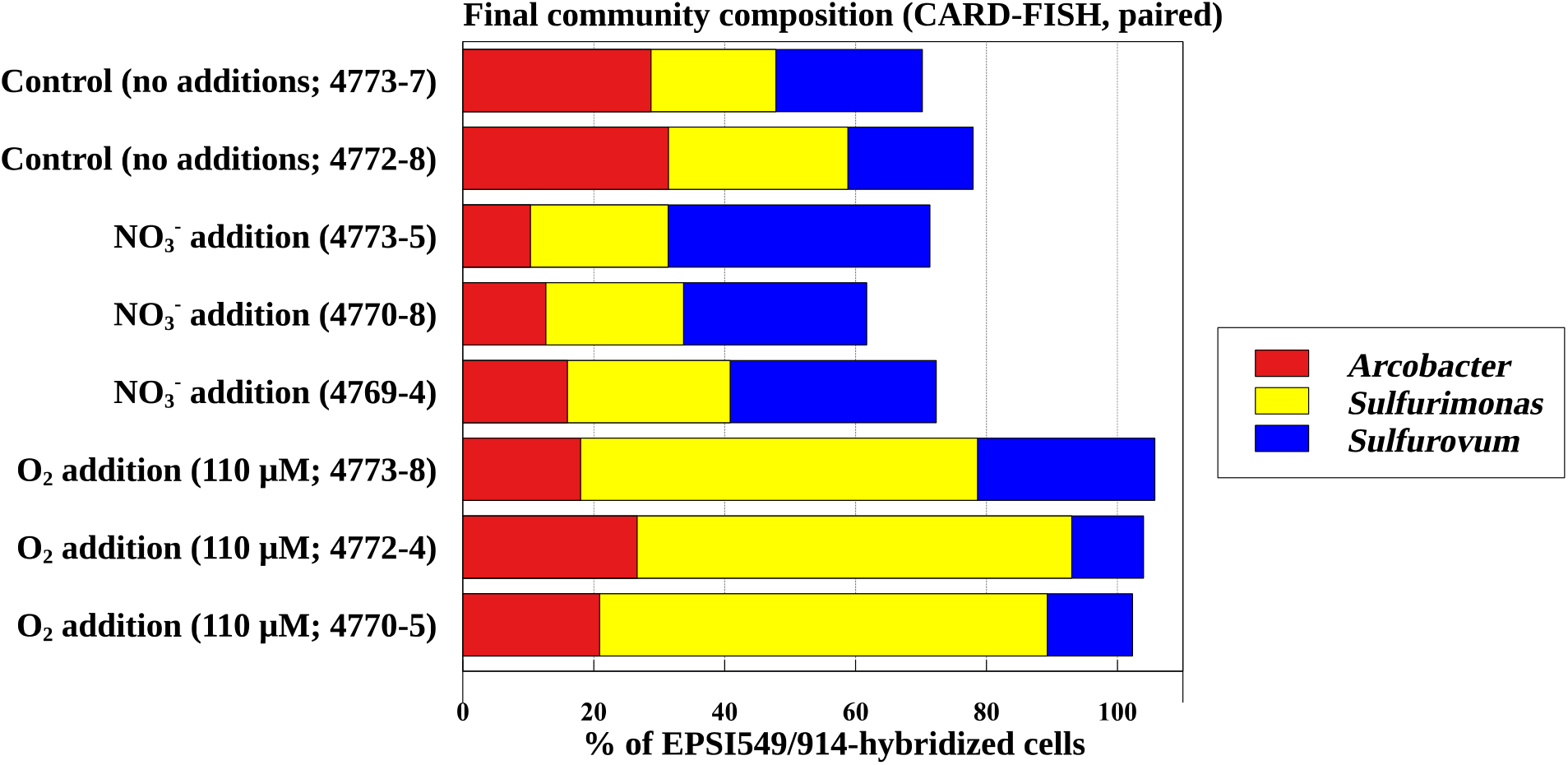
Campylobacterial community composition as determined by CARD-FISH at the end of *paired incubations* (i.e. non-radiocarbon incubations, corresponding to the chemical and DNA measurements).

**Figure S2:**
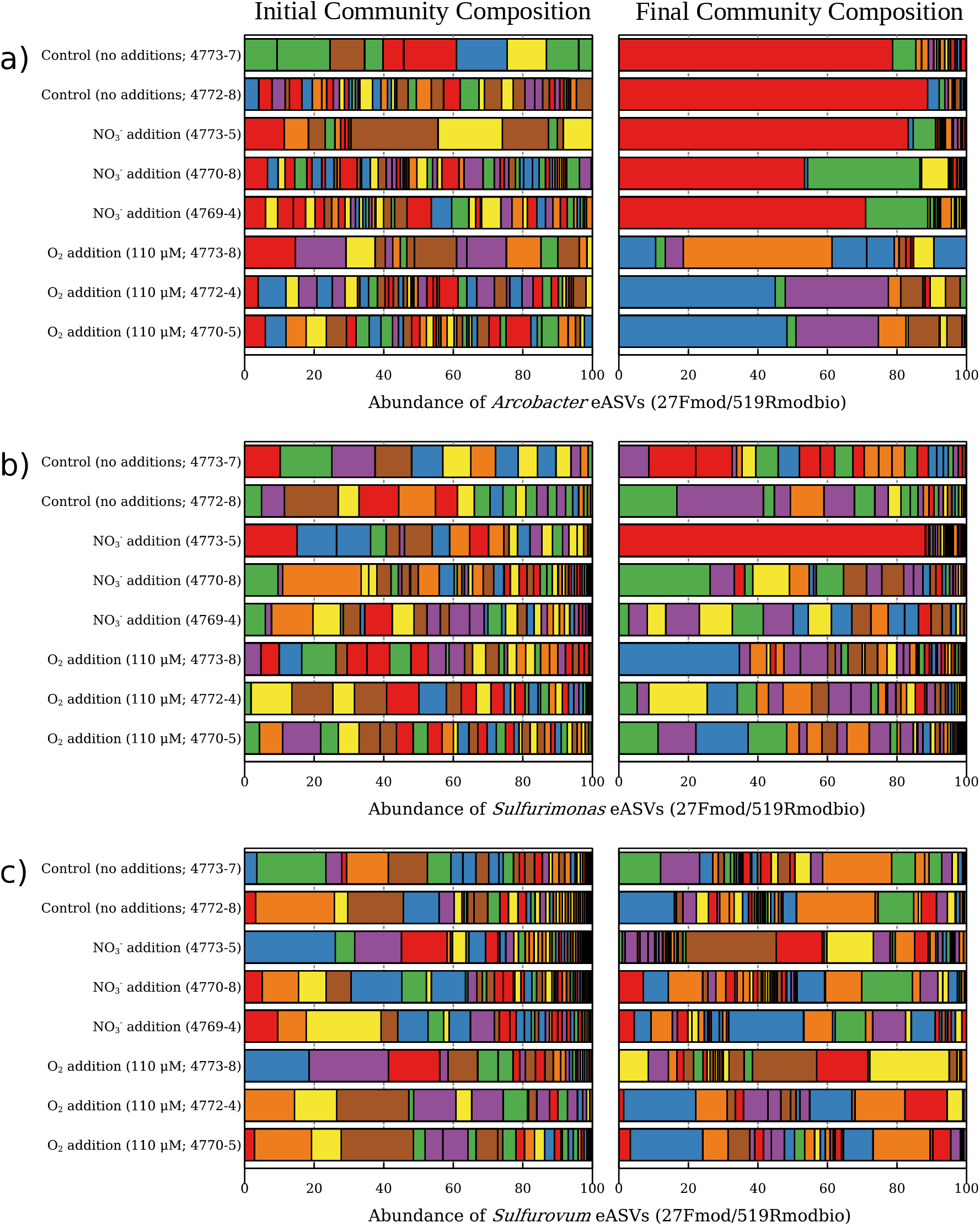
Comparisons of initial (left panel) and final (right panel) community composition from paired incubations as determined by 16S amplicon sequencing. The three panels show the community composition at the exact amplicon sequence variant (eASV) level for *Arcobacter*, *Sulfurimonas,* and *Sulfurovum*, respectively.

## Footnotes

1 Although operationally classified as genera, we use the term “group” to indicate the likelihood that these names - in particular *Sulfurovum* - represent broader phylogenetic groupings as noted by others (Meier et al 2017).

## Notes

https://osf.io/yts4p/

https://osf.io/9e6ra/

https://www.ncbi.nlm.nih.gov/bioproject/PRJNA505918

### Competing Interest Statement

The authors have declared no competing interest.

